# Effect of Melatonin on Tau aggregation and Tau-mediated cell surface morphology

**DOI:** 10.1101/861237

**Authors:** Rashmi Das, Abhishek Ankur Balmik, Subashchandrabose Chinnathambi

**Affiliations:** Neurobiology Group, Division of Biochemical Sciences, CSIR-National Chemical Laboratory, Dr. Homi Bhabha Road, 411008 Pune, India; Academy of Scientific and Innovative Research (AcSIR), 110025 New Delhi, India

**Keywords:** Alzheimer’s disease, Tau protein, Paired helical filaments, Tauopathies, Melatonin

## Abstract

Tau is the major neuronal protein involved in the stabilization of microtubule assembly. In Alzheimer’s disease, Tau self assembles to form intracellular protein aggregates, which are toxic to cells. Various methods have been tried and tested to restrain the aggregation of Tau. Most of the agents tested for this purpose have limitations in their effectiveness and availability to neuronal cells. We tested melatonin against *in vitro* Tau aggregation and observed its effect on membrane topology, tubulin network and Tau phosphorylation in neuro2a and N9 cell lines. The aggregation and conformation of Tau was determined by ThT fluorescence and CD spectroscopy respectively. The morphology of Tau aggregates in presence and absence of melatonin was studied by transmission electron microscopy. Melatonin was found to reduce the formation of higher order oligomeric structures without affecting the overall aggregation kinetics of Tau. Melatonin also modulates and helps to maintain membrane topology as evidenced by FE-SEM analysis. Overall, melatonin administration shows mild anti-aggregation and cytoprotective effects.

## Introduction

Microtubule-associated protein Tau undergoes misfolding and forms intracellular aggregates in Alzheimer’s disease (AD) and related Tauopathies ^*1*^. Deposition of extracellular fibrillar aggregates of amyloid-β also has deleterious effects on neuronal functions ^*2, 3*^. Both the proteinaceous aggregates have been widely studied with respect to their pathological consequences in AD and form the basis of two hypotheses known as ‘Tau’ and ‘amyloid-β’ hypothesis. Tau hypothesis has attained more attention in last decade since only amyloid-β deposition are found in brain without any pathological effect ^*4*^. The main physiological functions of Tau include the stabilization of microtubule, axonal growth and cargo trafficking *etc*. Under pathological condition, a series of signalling cascade leads to phosphorylation of Tau by various kinases. Tau is hyperphosphorylated upon upregulation of several kinases like CDK5 and GSK-3β. Therapeutic strategies have been employed involving the use of specific kinase inhibitors^*5-7*^. CDK5 complexed with p25 increases its activation state and induces the phosphorylation of Tau at the following amino acids Ser202, Thr205, Ser235 and Ser404. Additionally, the amyloidogenic protein aggregates accumulate around the phospho-lipid bilayer which leads to the formation of membrane pore in cholesterol-rich domain ^*8, 9*^. AD is associated with a multiple factors, which ultimately lead to neurodegeneration. Thus, multi-targeted approach is required for the effective amelioration of Tau-mediated neuropathology ^*10*^. Therapeutic strategies have been studied involving different aspects of neurodegeneration. A wide range of small molecules with multiple targets like aggregation inhibition, kinase inhibition or phosphatase activation are under study ^*7*^. Melatonin is a neurohormone-derived from pineal gland and involved in the maintenance of circadian rhythm ^*11, 12*^. Melatonin is also produced in non-pineal sites and its receptors are present in most cell types ^*13, 14*^. It is rapidly metabolized in liver and converted to secondary metabolites called kynuramines^*15-17*^. It is well known that melatonin is an anti-oxidant molecule, which can cross all biological membranes and also functions in the upregulation of other enzymes involved in protection against oxidative damage^*18-23*^. Melatonin is a pleotropic molecule with a high bioavailability and low toxicity. It is considered as a potent candidate against neurodegenerative diseases due to its wide range of functions. The role of melatonin in kinase function has been studied where it was found to downregulate kinases like GSK-3β and PKA, which in turn affects the phosphorylation status of Tau^*24-27*^. Melatonin is found to be effective in amelioration of neurological defects and restoration of cognitive functions ^*28, 29*^. Melatonin has also been studied with respect to inhibition of amyloid-β and α-synuclein aggregation^*30, 31*^. The multi-faceted role of melatonin makes it a potent lead for therapeutic approach ^*32*^.

Here, we have studied the potential role of melatonin in inhibition of full-length Tau (4R2N) aggregates and its effect on cell viability. We have also performed the function of melatonin in CDK5-mediated phosphorylation of Tau (Ser202/Thr205: AT8) sites. Melatonin reduces the membrane roughness induced by Tau aggregates in neuro2A and N9 microglial cells while the tubulin network remain unaltered by melatonin and Tau aggregates.

## Materials and methods

### Materials

Melatonin, BES, MES, Glycine, SDS, ThT, Protease inhibitor cocktail and ANS were purchased from Sigma-Aldrich. Other biochemical or molecular biology grade chemicals used were - DTT and IPTG (Calbiochem), APS, NaCl, PMSF, Sodium Azide, Ampicillin (MP Biomedicals), Luria-Bertani broth (Himedia), Acrylamide, EGTA, TEMED, DMSO (Invitrogen). BCA assay reagents used for protein estimation were purchased from Sigma-Aldrich,. Filtration devices were used from Merck and Pall Life sciences. Ethanol (Mol Bio grade), Chloroform, Isopropanol (Mol Bio grade) were purchased from MP Biomedicals; For cell culture studies, Dulbecco modified eagle’s media (DMEM), Fetal bovine Serum (FBS), Horse serum, Phosphate buffer saline (PBS, cell biology grade), trypsin-EDTA, Penicillin-streptomycin, RIPA buffer were also purchased from Invitrogen. MTT reagent, Okadaic acid, Paraformaldehyde (16%), TritonX-100 were purchased from Sigma. The coverslip of 0.17 mm was purchased from blue star for immunofluorescence and SEM study. In immunofluorescence and western blot study we used the following antibodies: mouse Beta Tubulin loading control (BT7R) (Thermo fisher, cat no MA516308), mouse monoclonal CDK5 antibody (invitrogen, Cat no AHZ0492) and total Tau antibody K9JA (Dako, Cat no A0024), AT8 (Thermo fisher, Cat no MN1020), Goat anti-Rabbit IgG (H+L) Cross-Adsorbed Secondary Antibody HRP (Invitrogen, A16110), anti-mouse secondary antibody conjugated with Alexa flour-488 (Invitrogen, Cat no A-11001), Goat anti-Rabbit IgG (H+L) Cross-Adsorbed Secondary Antibody with Alexa Fluor 555 (A-21428), DAPI (Invitrogen). For real-time quantification of cytokine expression profile, Trizol reagent, First strand cDNA synthesis kit (Cat no K1612), Maxima SYBR Green/Fluorescein qPCR Master Mix (2X) (Cat no K0241) were used from Thermo.

### Protein expression and Purification

pT7C full-length Tau (hTau40wt) were expressed in *E.coli* BL21* upon induction by 0.5 mM IPTG and purified as previously described ^*33-35*^. Cell homogenization was carried out at 15 KPSI using Constant cell disruption system. 0.5 M NaCl and 5 mM DTT were added to the resulting lysate and kept in water bath at 90°C for 20 minutes. The lysate was cooled, centrifuged at 40000 rpm for 45 minutes and dialyzed overnight. After a second round of centrifugation, the supernatant was filtered and subject to cation exchange chromatography using Sepharose fast flow (SPFF) column pre-equilibrated with 20 mM MES pH 6.8, 50 mM NaCl. For eluting Tau protein, 20 mM MES pH 6.8 with 1 M NaCl was used. The fractions containing Tau protein were pooled, concentrated and subjected to size-exclusion chromatography using 1X PBS buffer with 2 mM DTT in Superdex 75 Hi-load 16/600 column. The final concentration of Tau was obtained by performing Bicinchoninic acid assay.

### Aggregation inhibition assay

Full-length Tau aggregates were prepared in 20 mM BES pH 7.4 as assembly buffer, in the presence of heparin as an inducer (Tau: heparin= 4:1)^*36*^. The reaction was set-up in microcentrifuge amber tubes and incubated at 37°C for 96 hours. Aggregation reaction tubes with 200-5000 µM of melatonin in 10% DMSO were also prepared. ThT fluorescence for all reactions at various time-points (0-96 hours) was measured at 450 nm excitation and 475 nm emission wavelengths. The measurements for all time-points were taken in triplicates with Tau and ThT ratio of 1:2. Fluorescence of assembly buffer was measured for blank subtraction.

### CD spectroscopy

CD spectra for 3 µM soluble Tau and Tau aggregates with or without melatonin were recorded in Jasco J-815 CD spectrometer in 1 mm cuvette under nitrogen atmosphere. The spectra were recorded at 100 nm/min scan speed at 1 nm bandwidth in 190-250 nm scan range. The baseline was recorded for 50 mM sodium phosphate buffer pH-6.8 and subtracted for each spectrum. The final spectra were obtained as an average of five CD acquisitions.

### Transmission Electron Microscopy

Morphology of Tau aggregates with or without melatonin were visualized by transmission electron microscopy. 2 µM of Tau aggregates were placed on 400 mesh carbon coated copper grids, washed twice with filtered miliQ water for 30 seconds and stained with 2% uranyl acetate for 2 minutes. The samples were scanned using TECNAI T20 Transmission Electron Microscope at 120 KV.

### Size-exclusion chromatography

Full-length Tau aggregates were prepared in 20 µM BES, pH 7.4 in 4:1 (Tau: heparin) ratio with heparin in the presence of 1000 µM melatonin and additives as mentioned earlier. A control experiment was performed with only full-length Tau aggregates. Size-exclusion chromatography was performed at 0 hour after 24 hours using 1X PBS buffer for elution. The respective samples were loaded onto Superdex Increase 10/300 GL column in 100 µl sample volume after filtration through 0.45 µm centrifugal filters. The peak fractions of 0.25 ml were collected for each run.

### Cytotoxicity assay for SEC fractions

The fractions obtained from size-exclusion chromatography of full-length Tau aggregates in presence or absence of 1000 µM melatonin were tested for their effect on viability of neuro2A cells (Fractions F7-F10 corresponds to Tau with melatonin, Fractions F11-F14 corresponds to Tau only from SEC). Neuro2A cells were cultured in DMEM with 10% FBS and 100µg/mL of penicillin and streptomycin. Neuro2A cells were seeded in 96 well plates (10000 cells/well) and incubated at 37°C for 12 hours at 5% CO2. Peak fractions (F7-F10) were obtained from SEC of full length Tau with melatonin and fractions (F11-F14) were obtained from SEC of Tau aggregates. Cells were treated with 30 µl of fractions and 70 µl of reduced serum media (DMEM with 0.5% FBS) in each well and incubated for 24 hours. MTT assay was performed to check the cell viability. MTT was added at 0.5 mg/ml concentration in each well and incubated for 3 hours. The formazan end product solubilized by adding absolute DMSO. The spectrophotometric absorbance was measured at 570 nm in triplicate (Tecan Infinite 200 Pro).

### Real-time PCR

Neuro2A cells were treated with melatonin (50 µM) and okadaic acid (25 nM) for 6 hours. Then the cells were washed in PBS and total RNA were isolated by the conventional TRIZOL, chloroform and isopropanol procedure. The RNA pellets were proceeded for cDNA synthesis (First-strand cDNA synthesis kit) with oligo-dT primer. The expression level of CDK5 was checked by quantitative PCR. The fold change was calculated by ΔΔCT method with respect to house-keeping GAPDH control (Table 1).

### Western Blot

Neuro2A cells with okadaic acid (25 nM) and melatonin (50 µM) separately and together for 24 hours. The cells were washed with PBS, lysed with RIPA buffer and cell lysate were subjected to western blot with anti-CDK5 monoclonal antibody (1:1500) with β-tubulin (1:5000). Then, the bands intensity was quantified by using BIORAD Quality one 4.6.6 software. The band-density of the treated groups were compared with its corresponding untreated control group and the relative fold changes were plotted with respective proteins and treated group.

### Immunofluorescence microscopy

Misfolded Tau becomes hyperphosphorylated in case of AD due to over-activation of various protein kinases. Phosphorylated Tau tends to aggregate inside the cells and subsequently secreted outside which lead to improper functioning. To study the over-activation of one of the cellular kinases-CDK5 and intracellular Tau phosphorylation, we treated the neuro2A cells with okadaic acid (25 nM) and melatonin (50 µM) separately and together for 24 hours. The cells were washed with PBS thrice and fixed with 4% paraformaldehyde for 15 minutes and permeabilized with 0.2% TritonX-100. The cells were stained with AT8 (1:100), K9JA (1:500), CDK5 (1:50) and mouse anti-tubulin antibody (1:100) for overnight in 2% horse serum-PBS. Then, Alexa flour conjugated secondary antibody was allowed to bind p-Tau, CDK5 specific primary antibody (anti-mouse Alexa flour 555, 1:500) and K9JA (anti-rabbit Alexa flour 488, 1:1000) for 1 hour along with nuclear staining DAPI (300 nM) for 5 minutes. The images and 3D localization (orthogonal view) were observed in Zeiss Axio observer with Apotome2 fluorescence microscope at 63X oil immersion and analyzed by ZEN2 software.

### Surface morphology by Field-Emission Scanning electron microscopy (FE-SEM)

To study the surface morphology of neuro2A and microglia (N9) cells by melatonin and Tau aggregates treatment, the cells were grown onto coverslip for 16 hours at 37°C, 5% CO_2_. Then the cells were washed with PBS twice and treated individually and in combination with 10 µM of Tau aggregates and 50 µM of melatonin for 6 hours. After the treatment, cells were fixed with 2.5% glutaraldehyde for 1 hour at 4°C, followed by step-wise dehydration with 10, 25, 50, 75, 95 and 100% ethanol. The cells were kept overnight in vacuumed desiccator in presence of anhydrous calcium carbonate for complete drying. The complete coverslips were coated with gold nanoparticle and scanned in FE-SEM (FEI- NOVA NANOSEM 450) at 8000 or 10000X magnification with 18 KV electron beam.

### Statistical analysis

All the experimental analyses were carried out in triplicate. The statistical analysis of control and treated samples were carried out by SigmaPlot 10.0 (Systat software). Two-tailed student unpaired t-test was performed to show the significance. (n.s.-non-significant, ** indicates P ≤ 0.01, *** indicates P ≤ 0.005).

## Results and discussion

### Melatonin shows no effect on aggregation of full-length Tau

The effect of melatonin in the inhibition of full-length Tau protein (Fig. 1A) aggregation was studied using ThT fluorescence assay. Melatonin was used against Tau aggregation at a concentration range of 200-5000 µM. Surprisingly, it did not show any significant effect on the aggregation of full-length Tau protein in ThT fluorescence, even with higher concentration (5000 µM) of melatonin (Fig. 2A). Also, CD spectroscopy revealed that melatonin treatment does not bring about significant structural change. Tau aggregates shows the signature for β-sheet structure, which remained unchanged with melatonin treatment (Fig. 1C). However, the electron microscopy images for full-length Tau aggregation samples in presence of melatonin at 1000 and 5000 µM showed distinct morphology of Tau fibrils where mostly broken filaments were observed (Fig. 1D). Melatonin has been studied earlier in the inhibition of amyloid-β and α-synuclein aggregates ^*30*^. The mode of action of melatonin involves the disruption of salt-bridge formation or reducing the hydrophobic interaction between proteins^*31, 37*^. For Tau aggregation inhibition, melatonin may employs the same mechanism but it is required in higher concentration to disrupt the fibril formation.

**Figure 1.**
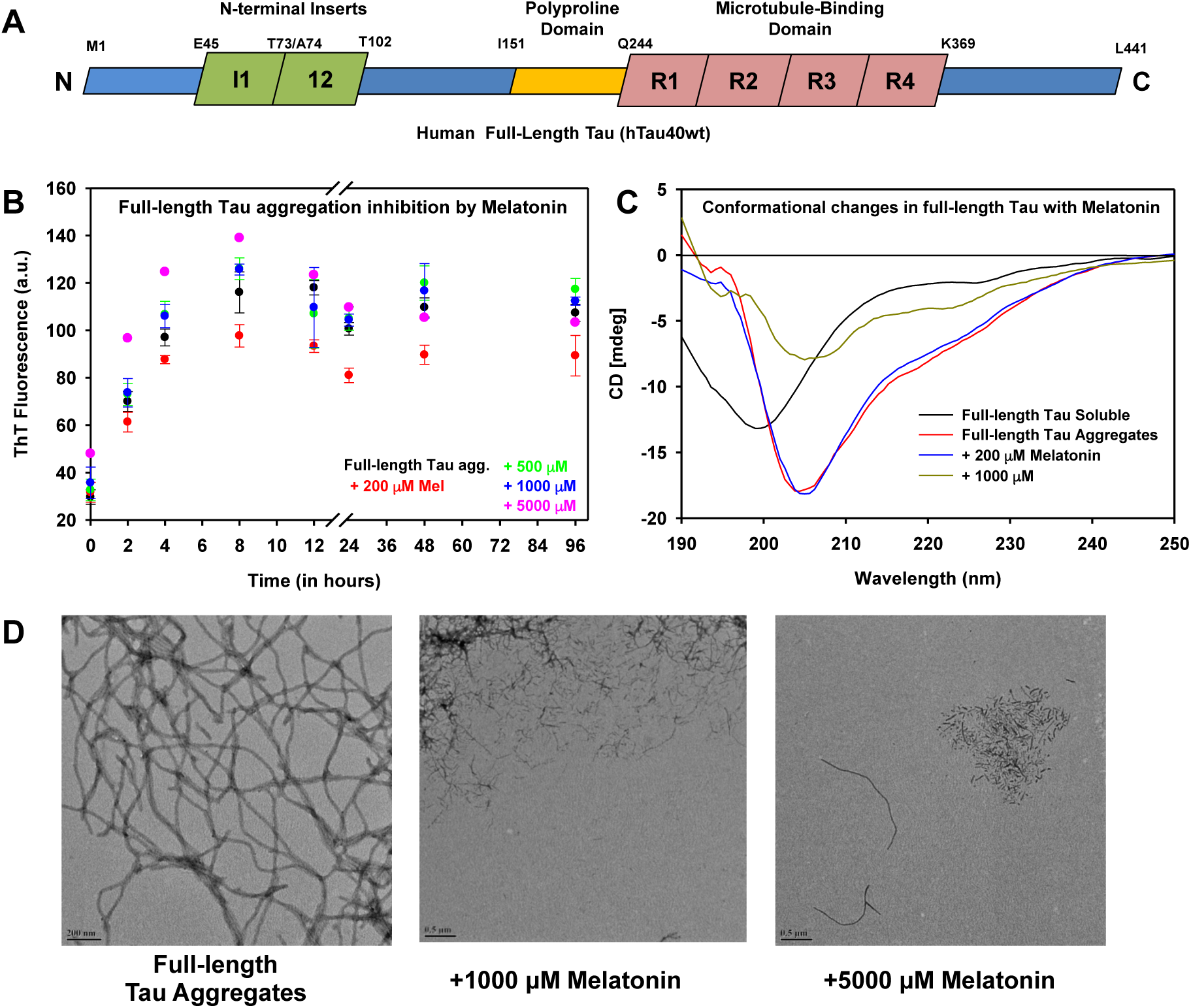
Effect of melatonin on full-length Tau aggregation. A) Domain organization of human full-length Tau consists of two N-terminal inserts and four imperfect repeat domains. The four repeat domains hold importance as the microtubule-binding domain of Tau as well as the core of the pathological PHF assembly. The polyproline domain plays an important role in the phosphorylation of Tau *via* proline-directed kinases. B) Aggregation inhibition assay for full-length Tau monitored by ThT fluorescence. There was no change in ThT fluorescence in samples incubated with melatonin in as high concentration as 5000 µM. C) CD spectroscopy for full-length Tau incubated with melatonin at 200 and 1000 µM concentration. CD spectrum for Tau control aggregates was also measured. At higher concentrations of melatonin, spectra shift towards random coil. There are no conformation changes in full-length Tau sample with 200 µM of melatonin as compared to control Tau aggregates. D) TEM images for full-length Tau control aggregates showed mature fibrils while in melatonin treated samples (1000 and 5000 µM) broken aggregates were observed.

**Figure 2.**
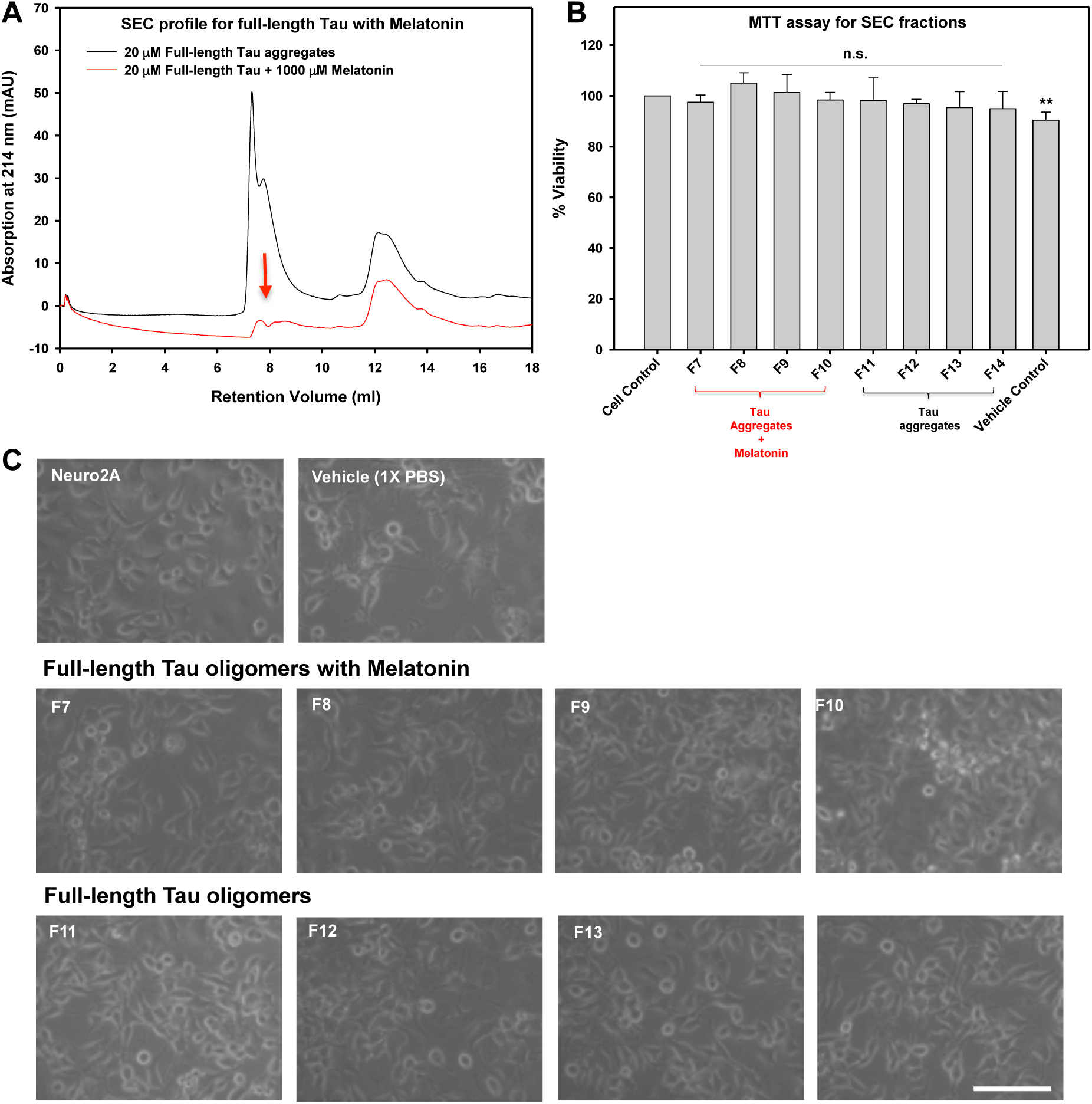
Size-exclusion chromatography for full-length Tau upon melatonin treatment. A) Size-exclusion chromatography for 20 µM hTau40wt aggregates with 1000 µM melatonin as compared to hTau40wt aggregates control after 24 hours of incubation. The higher order oligomer peak for Tau disappears in the sample incubated with melatonin. B) Cell viability assay for the fractions obtained from SEC runs for hTau40wt aggregates control and hTau40wt aggregates with melatonin. No toxicity was observed from the fractions obtained from either SEC (n.s.-non-significant, ** indicates P ≤ 0.01, *** indicates P ≤ 0.005). C) Bright field images for neuronal cells treated with SEC fractions showed morphology similar to untreated cells (Scale bar – 100 µm).

### Melatonin reduces the formation of higher order oligomeric species of Tau

The aggregation of Tau protein forms intermediate toxic oligomers, which also act like seeding species for further aggregation ^*38*^. We prepared Tau aggregates in the presence or absence of melatonin and carried out SEC for the separation of oligomeric species ^*39*^. For the preparation of oligomers, full-length Tau was incubated at 37 °C in the presence of heparin with or without melatonin. Size-exclusion chromatography (SEC) was performed after 24 hours to separate these oligomers and to see its effect on viability of neuro2a cells upon exposure. When the aggregates were prepared in presence of melatonin, the higher order oligomer were not observed in SEC. In SEC experiment, the peak for full length Tau oligomeric species of lower order was obtained as suggested by the retention volume. These oligomeric species were absent in SEC of full-length Tau aggregates in the presence of melatonin (indicated by downward red arrow) (Fig. 2A). The fractions collected from SEC for full-length Tau aggregates and corresponding fractions from SEC for Tau aggregates with melatonin showed no cytotoxicity when treated to neuro2A cells. It also had no effect on morphology of cells after treatment for 24 hours (Fig 2C) probably due to the lower oligomer concentration or non-toxic forms of oligomer species. The results showed that melatonin can effectively inhibit the formation of these oligomeric species.

### Phospho Tau (AT8) and CDK5 levels remained unaltered by melatonin

Misfolded Tau can undergo several post-translational modifications, among which phosphorylation is the most frequent (80 possible serine, threonine, tyrosine residues) ^*1*^ which consecutively induces Tau aggregation ^*40*^. CDK5 complexed with p25 (CDK5-p25), with enhanced kinase activity which induces hyperphosphorylation of Tau in AD (at Threonine 231, Serine 202/Threonine 205, Serine 262, Serine 396) ^*41*^. Melatonin only maintained the basal level expression of CDK5 level on neuro2A cells as observed by qRT-PCR and western blot (Fig. 3A, B, C) while OA-induced the CDK5 level in melatonin-treated neuro2A cells as seen by immunostaining (Fig. 3D). CDK5 level remains unaltered at different treatment group as observed by fluorescence microscopy (Fig. 3E).

**Figure 3.**
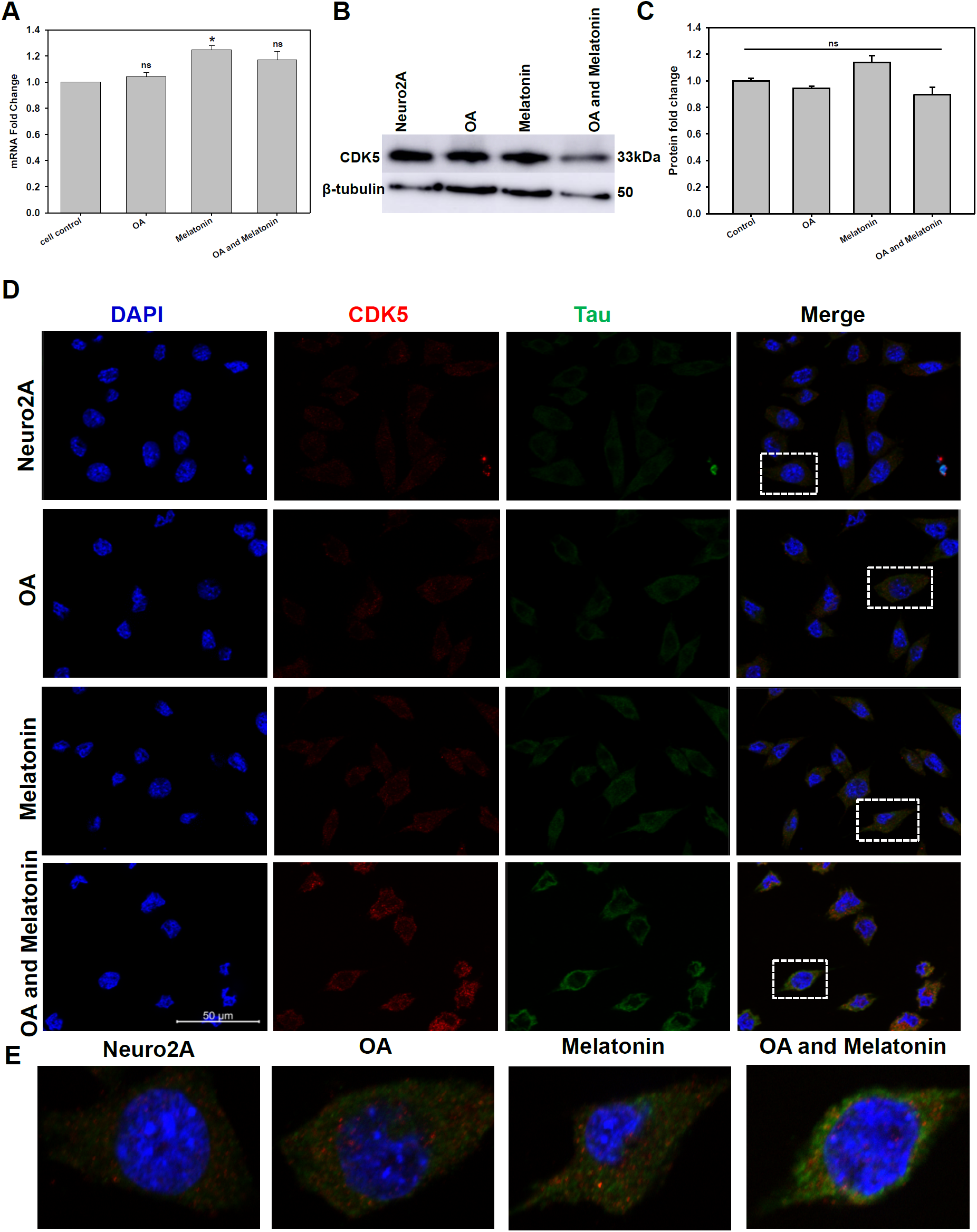
Cellular level of CDK5 and Tau upon OA and melatonin exposure by immunofluorescence study. (A, B, C) CDK5 mRNA expression level and protein level remain constant in neuro2a cells upon the exposure of melatonin and OA treatment. (D) The CDK5 and Tau (K9JA) level remain constant in neuro2A cells with melatonin and OA treatment. But combined treatment with both has increased the CDK5 level slightly. (E) Enlarged single cell images were shown for different treatment groups.

To confirm the localization of p-Tau into the neuro2A cells upon same treatment, cells were immunostained with AT8 antibody along with total-Tau K9JA antibody as a control. Neuroblastoma cells have expressed a constant level of intracellular AT8 epitope of Tau (Fig. 4A) whereas; OA and melatonin treatment did not alter basal level phospho-Tau expression inside the nucleus (Fig. 4B). In our study, we have showed the constant level of CDK5 and AT8-Tau throughout the groups. This signifies that okadaic acid as well as melatonin did not involve in CDK5-mediated Tau phosphorylation.

**Figure 4.**
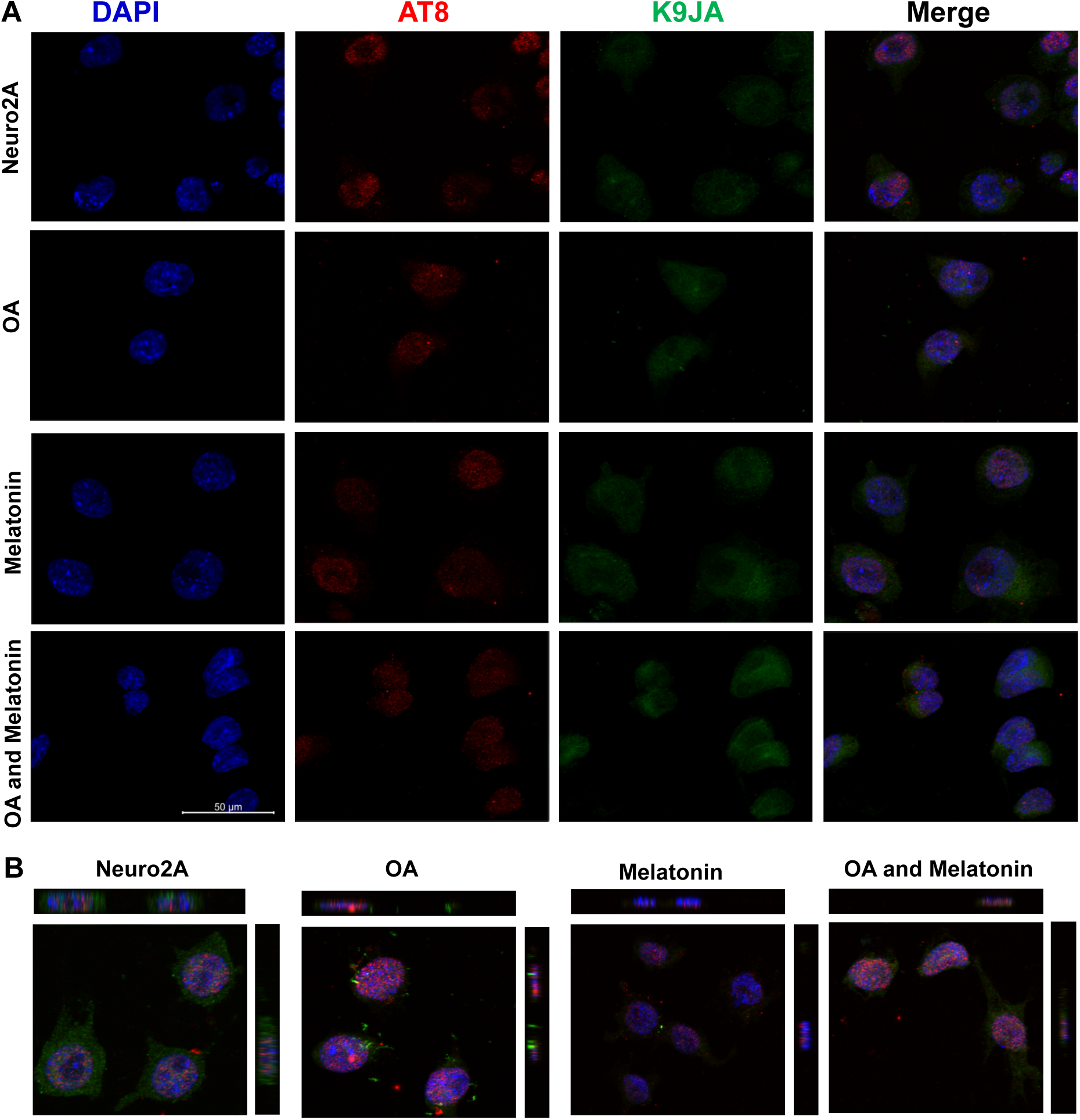
Level of Phospho-Tau (AT8) and total Tau (K9JA) in OA and melatonin treatment by immunofluorescence study. (A) OA treatment did not altered the level of Tau phosphorylation (AT8: S202/ T205) on neuro2A cells while melatonin also did not alter the level of AT8 immunostaining, localized mainly in nucleus. (B) Orthogonal projection for 3D localization of AT8-Tau inside the nucleus while unmodified Tau was dispersed throughout the cytosol.

### Alteration of membrane morphology and Tubulin network by Tau aggregates and melatonin

The plasma membrane topography is a determinant of normal physiological or pathological state of neuro-glial cells. The alteration of membrane structure occurs either due to changes in raft region or the variation of cytoskeleton configuration. The membrane topography can be broadly two types: roughness and stiffness. Aβ which is one of the key protein responsible for AD, increases membrane stiffness ^*42*^ whereas; Taxol reduces membrane roughness by stabilizing microtubules ^*43*^. Neuro-glial cells when treated with Tau aggregates, it induced membrane roughness as compared to without treatment control cells (Fig. 5A). The membrane roughness was more in case of neuro2A cells (Fig. 5A) whereas; bulging type structures were more prominent in N9 cells (Fig. 6A). As it is well proved that Tau is a microtubule stabilizing protein and Tau aggregation leads to microtubule disorganization during AD, extracellular Tau aggregates may influence the cytoskeleton mis-orientation as found by FE-SEM images. Similarly, microglia are important in spreading of Tau by exosomal release ^*44*^ and astrocytes become reactivated by microglial cross-talk ^*45*^. External Tau aggregates may induce microtubule disorganization, or membrane raft dislodging and improper vesicular release in this immune cell and neuronal cells. Melatonin reduced the degree of membrane roughness after Tau aggregates treatment, which may have a role in cytoskeleton stabilization and membrane raft organization (Fig. 5, 6A).

**Figure 5.**
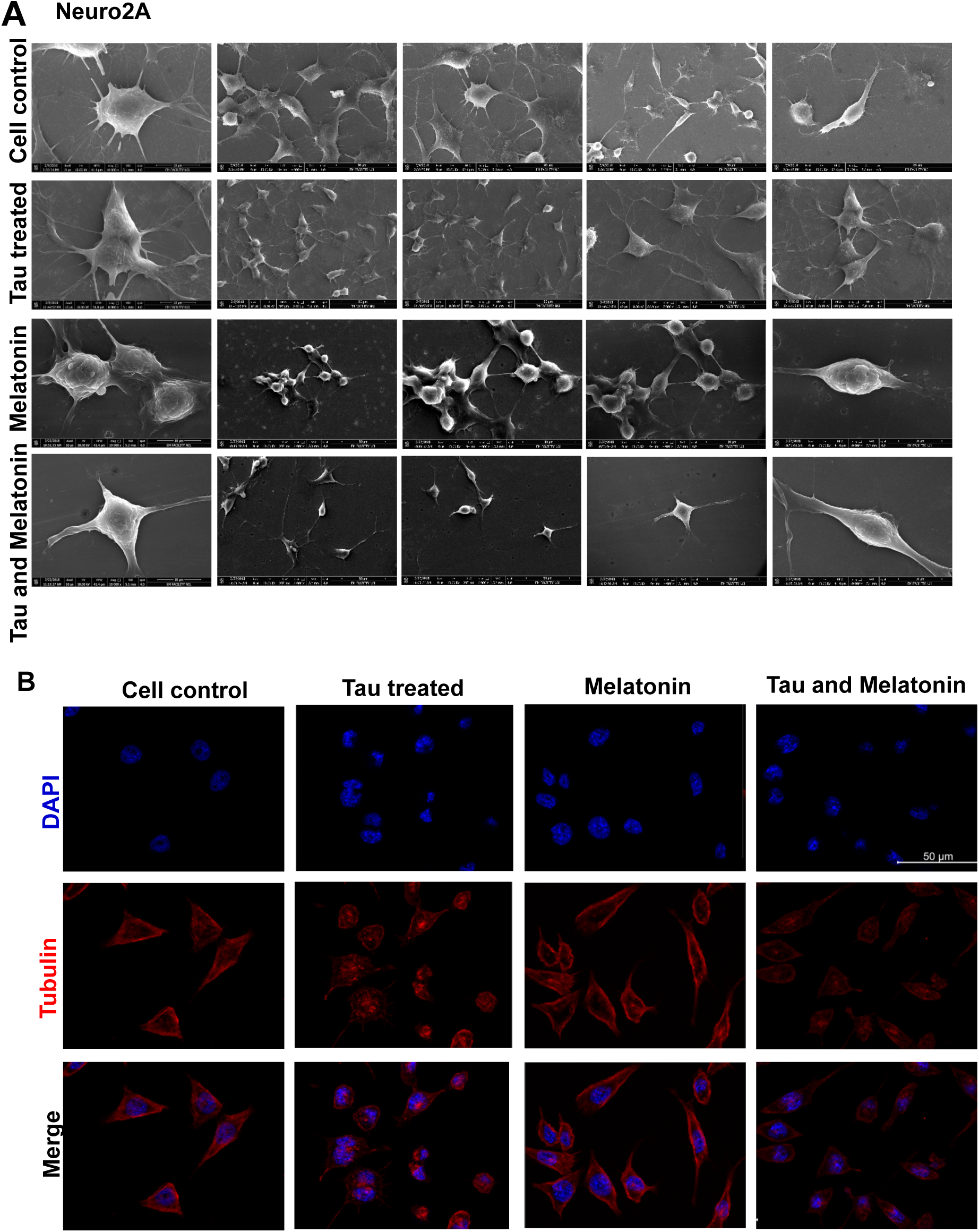
Alteration of membrane morphology of neuro2A and tubulin network by FE-SEM and immunofluorescence study. (A) Untreated control revealed smooth membrane morphology of neuro2A. Tau aggregates treatment showed membrane roughness on neuro2A cells while melatonin treatment restored the membrane stiffness completely on neuro2A cells. Melatonin showed unaltered membrane morphology on neuro2a, (B) Tau aggregates and melatonin treatment was showed no alteration in intracellular tubulin network into the neurons as compared to untreated control.

**Figure 6.**
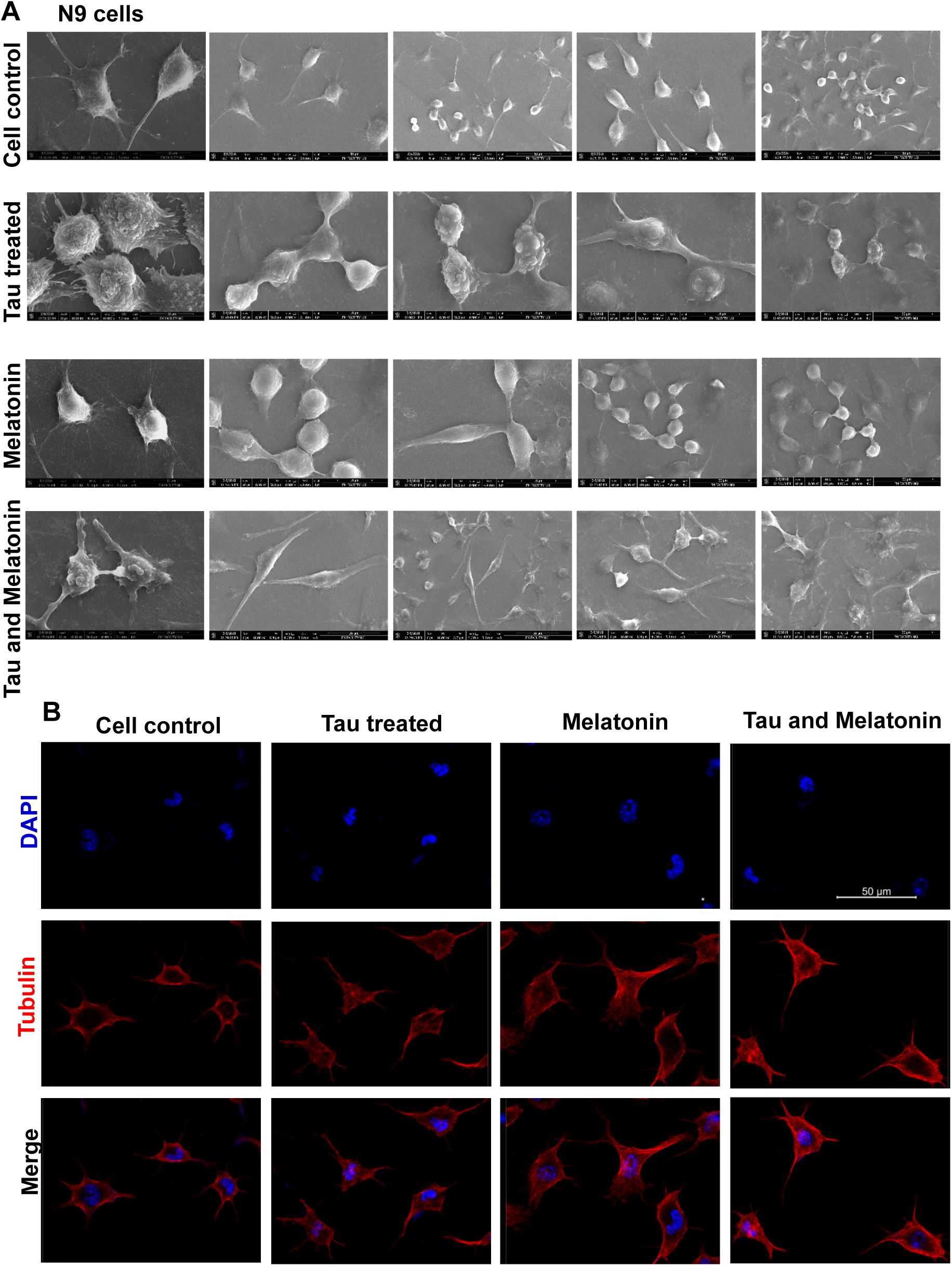
Missorting of tubulin Network and membrane morphology in Tau aggregates and melatonin-treated microglial cell (N9). (A) Tau aggregates treatment showed membrane blebbing on N9 cells while melatonin treatment showed restored membrane morphology. (B) Tau aggregates treated N9 cells showed extended morphology and melatonin treatment rearranged the tubulin throughout the cell.

Microtubules stabilize the cell shape, structure and facilitate several plasmalemma events-like migration, exosomal release, endocytosis, induction of signalling cascade *etc*. To study the effect of Tau aggregates and melatonin on cytoskeletal network, cellular tubulin was immune-stained after the same treatments as mentioned above. Extracellular Tau aggregates-induced unaltered tubulin patches in the cytoplasm whereas; melatonin treatment showed intense tubulin network in neuro2A cells (Fig. 5B). Tau aggregates treatment might activate the microglia with more extended cell bodies compared to untreated control. Melatonin treatment did not alter the microtubule network in microglia (Fig. 6B). In our study, we evidenced the initiation of membrane roughness on neurons and microglial cells upon Tau fibrils exposure. But the event was reinstated by melatonin which signifies its role in remodelling of membrane dynamics and cell-to-cell communication in neurodegenerative diseases.

### Conclusion

Melatonin is a neurohormone, which serves a wide variety of function apart from maintenance of circadian rhythm. It mediates its cytoprotective action as an anti-oxidant itself as well as regulator of other proteins involved in anti-oxidant and anti-inflammatory function. Our studies shows *in vitro* administration of melatonin does not affect the aggregation of full-length Tau but it does reduce higher order oligomers. Melatonin also affected Tau phosphorylation and helps to maintain tubulin network and membrane topology in neuro2a and N9 cells.

## Abbreviations used

NFTs: neurofibrillary tangles,
PHF: paired helical filaments,
Aβ: amyloid-β,
CDK5/p25: cyclin dependent kinase 5/p25.
GSK-3β: Glycogen synthase kinase - 3β.

## AUTHOR INFORMATION

### Author Contributions

R.D., A.B. and S.C. designed the experiments. R.D. and A.B. carried out the experiments. R.D., A.B. and S.C. analyzed the data and wrote the article. S.C. conceived, supervised, resource produced and wrote the paper. All authors contributed to the discussions and manuscript review.

### Notes

The authors declare no competing financial interest.

## Acknowledgements

Rashmi Das acknowledges the fellowship from University Grant Commission (UGC) India. Abhishek Ankur Balmik acknowledges the Shyama Prasad Mukherjee fellowship (SPMF) from Council of Scientific Industrial Research (CSIR), India.

---

TOC

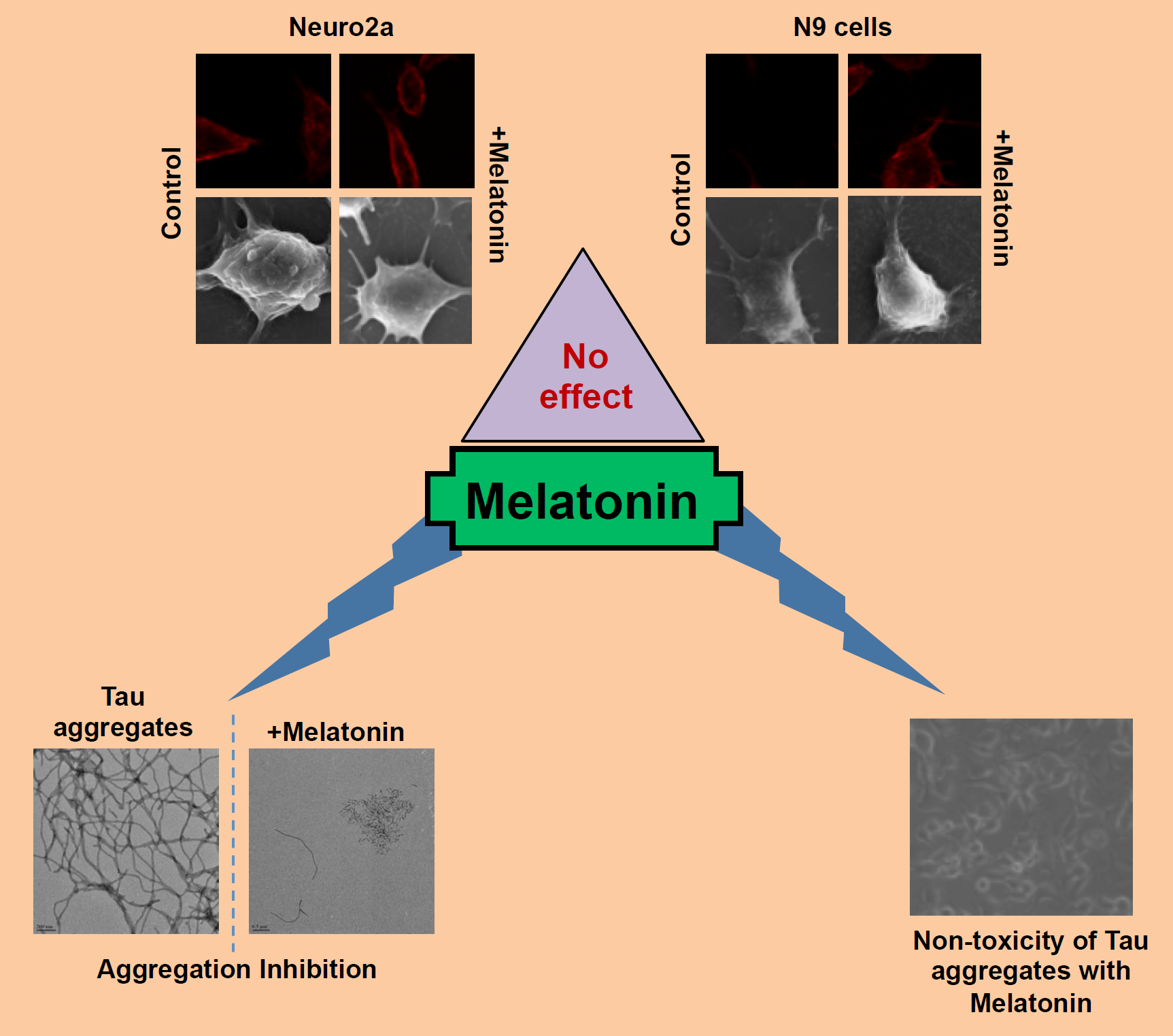

Modulation of membrane topology, tubulin network and Tau aggregation by melatonin

## References

1. Mandelkow, E.-M., and Mandelkow, E. (2012) Biochemistry and cell biology of tau protein in neurofibrillary degeneration, Cold Spring Harbor perspectives in medicine 2, a006247.

2. Dickson, D. W., and Yen, S.-H. C. (1989) Beta-amyloid deposition and paired helical filament formation: which histopathological feature is more significant in Alzheimer’s disease?, Neurobiology of aging 10, 402–404.

3. Hardy, J. A., and Higgins, G. A. (1992) Alzheimer’s disease: the amyloid cascade hypothesis, Science 256, 184.

4. Dickson, D. W., Crystal, H. A., Mattiace, L. A., Masur, D. M., Blau, A. D., Davies, P., Yen, S.-H., and Aronson, M. K. (1992) Identification of normal and pathological aging in prospectively studied nondemented elderly humans, Neurobiology of aging 13, 179–189.

5. Gong, C. X., and Iqbal, K. (2008) Hyperphosphorylation of microtubule-associated protein tau: a promising therapeutic target for Alzheimer disease, Current medicinal chemistry 15, 2321–2328.

6. Hilgeroth, A. P. D., and Tell, V. (2013) Recent developments of protein kinase inhibitors as potential AD therapeutics, Frontiers in cellular neuroscience 7, 189.

7. Congdon, E. E., and Sigurdsson, E. M. (2018) Tau-targeting therapies for Alzheimer disease, Nature Reviews Neurology, 1.

8. Künze, G., Barré, P., Scheidt, H. A., Thomas, L., Eliezer, D., and Huster, D. (2012) Binding of the three-repeat domain of tau to phospholipid membranes induces an aggregated-like state of the protein, Biochimica et Biophysica Acta (BBA)-Biomembranes 1818, 2302–2313.

9. McLaurin, J., and Chakrabartty, A. (1997) Characterization of the interactions of Alzheimer β-amyloid peptides with phospholipid membranes, European Journal of Biochemistry 245, 355–363.

10. Götz, J., Ittner, A., and Ittner, L. M. (2012) Tau targeted treatment strategies in Alzheimer’s disease, British journal of pharmacology 165, 1246–1259.

11. Claustrat, B., Brun, J., and Chazot, G. (2005) The basic physiology and pathophysiology of melatonin, Sleep medicine reviews 9, 11–24.

12. Pandi-Perumal, S. R., Trakht, I., Srinivasan, V., Spence, D. W., Maestroni, G. J. M., Zisapel, N., and Cardinali, D. P. (2008) Physiological effects of melatonin: role of melatonin receptors and signal transduction pathways, Progress in neurobiology 85, 335–353.

13. Stefulj, J., Hörtner, M., Ghosh, M., Schauenstein, K., Rinner, I., Wölfler, A., Semmler, J., and Liebmann, P. M. (2001) Gene expression of the key enzymes of melatonin synthesis in extrapineal tissues of the rat, Journal of pineal research 30, 243–247.

14. Reiter, R. J., and Tan, D.-X. (2003) What constitutes a physiological concentration of melatonin?, Journal of Pineal Research 34, 79–80.

15. Cajochen, C., Kräuchi, K., and Wirz-Justice, A. (2003) Role of melatonin in the regulation of human circadian rhythms and sleep, Journal of neuroendocrinology 15, 432–437.

16. Cardinali, D. P., Lynch, H. J., and Wurtman, R. J. (1972) Binding of melatonin to human and rat plasma proteins, Endocrinology 91, 1213–1218.

17. Hardeland, R., Tan, D. X., and Reiter, R. J. (2009) Kynuramines, metabolites of melatonin and other indoles: the resurrection of an almost forgotten class of biogenic amines, Journal of pineal research 47, 109–126.

18. Anton-Tay, F. D. J. L., Diaz, J. L., and Fernandez-Guardiola, A. (1971) On the effect of melatonin upon human brain. Its possible therapeutic implications, Life Sciences 10, 841–850.

19. Yu, H., Dickson, E. J., Jung, S.-R., Koh, D.-S., and Hille, B. (2016) High membrane permeability for melatonin, The Journal of general physiology 147, 63–76.

20. Tomas□Zapico, C., and Coto□Montes, A. (2005) A proposed mechanism to explain the stimulatory effect of melatonin on antioxidative enzymes, Journal of pineal research 39, 99–104.

21. Wang, X. (2009) The antiapoptotic activity of melatonin in neurodegenerative diseases, CNS neuroscience & therapeutics 15, 345–357.

22. Wang, J. Z., and Wang, Z. F. (2006) Role of melatonin in Alzheimer-like neurodegeneration, Acta Pharmacologica Sinica 27, 41–49.

23. Tan, D. X., Reiter, R. J., Manchester, L. C., Yan, M. T., El-Sawi, M., Sainz, R. M., Mayo, J. C., Kohen, R., Allegra, M. C., and Hardeland, R. (2002) Chemical and physical properties and potential mechanisms: melatonin as a broad spectrum antioxidant and free radical scavenger, Current topics in medicinal chemistry 2, 181–197.

24. Zempel, H., Thies, E., Mandelkow, E., and Mandelkow, E.-M. (2010) Aβ oligomers cause localized Ca2+ elevation, missorting of endogenous Tau into dendrites, Tau phosphorylation, and destruction of microtubules and spines., Journal of Neuroscience 30, 11938–11950.

25. Wang, D. L., Ling, Z. Q., Cao, F. Y., Zhu, L. Q., and Wang, J. Z. (2004) Melatonin attenuates isoproterenol-induced protein kinase A overactivation and tau hyperphosphorylation in rat brain, Journal of pineal research 37, 11–16.

26. Planel, E., Yasutake, K., Fujita, S. C., and Ishiguro, K. (2001) Inhibition of protein phosphatase 2A overrides Tau protein kinase I/glycogen synthase kinase 3b and cyclin-dependant kinase 5 inhibition and results in tau hyperphosphorylation in the hippocampus of starved mouse, Journal of Biological Chemistry.

27. Wang, J.-z., Wu, Q., Smith, A., Grundke-Iqbal, I., and Iqbal, K. (1998) τ is phosphorylated by GSK-3 at several sites found in Alzheimer disease and its biological activity markedly inhibited only after it is prephosphorylated by A-kinase., FEBS letters 436, 28–34.

28. Lin, L., Huang, Q.-X., Yang, S.-S., Chu, J., Wang, J.-Z., and Tian, Q. (2013) Melatonin in Alzheimer’s disease, International journal of molecular sciences 14, 14575–14593.

29. Zhou, J. N., Liu, R. Y., Kamphorst, W., Hofman, M. A., and Swaab, D. F. (2003) Early neuropathological Alzheimer’s changes in aged individuals are accompanied by decreased cerebrospinal fluid melatonin levels, Journal of pineal research 35, 125–130.

30. Ono, K., Mochizuki, H., Ikeda, T., Nihira, T., Takasaki, J.-i., Teplow, D. B., and Yamada, M. (2012) Effect of melatonin on α-synuclein self-assembly and cytotoxicity, Neurobiology of aging 33, 2172–2185.

31. Pappolla, M., Bozner, P., Soto, C., Shao, H., Robakis, N. K., Zagorski, M., Frangione, B., and Ghiso, J. (1998) Inhibition of Alzheimer β-fibrillogenesis by melatonin, Journal of Biological Chemistry 273, 7185–7188.

32. Balmik, A. A., and Chinnathambi, S. (2018) Multi-Faceted Role of Melatonin in Neuroprotection and Amelioration of Tau Aggregates in Alzheimer’s Disease, Journal of Alzheimer’s Disease 62, 1481–1493.

33. Gorantla, N. V., Shkumatov, A. V., and Chinnathambi, S. (2017) Conformational Dynamics of Intracellular Tau Protein Revealed by CD and SAXS. In Tau Protein pp 3–20, Springer.

34. Gorantla, N. V., Khandelwal, P., Poddar, P., and Chinnathambi, S. (2017) Global conformation of tau protein mapped by Raman spectroscopy. In Tau Protein pp 21–31, Springer, Humana Press.

35. Sonawane, S. K., Ahmad, A., and Chinnathambi, S. (2019) Protein-Capped Metal Nanoparticles Inhibit Tau Aggregation in Alzheimer’s Disease, ACS Omega 4, 12833–12840.

36. Gorantla, N. V., Das, R., Mulani, F. A., Thulasiram, H. V., and Chinnathambi, S. (2019) Neem Derivatives Inhibits Tau Aggregation, Journal of Alzheimer’s Disease Reports 3, 169–178.

37. Skribanek, Z., Baláspiri, L., and Mák, M. (2001) Interaction between synthetic amyloid-β-peptide (1–40) and its aggregation inhibitors studied by electrospray ionization mass spectrometry., Journal of mass spectrometry 36, 1226–1229.

38. Ward, S. M., Himmelstein, D. S., Lancia, J. K., and Binder, L. I. (2012) Tau oligomers and tau toxicity in neurodegenerative disease, Portland Press Limited.

39. Flach, K., Hilbrich, I., Schiffmann, A., Gärtner, U., Krüger, M., Leonhardt, M., Waschipky, H., Wick, L., Arendt, T., and Holzer, M. (2012) Tau oligomers impair artificial membrane integrity and cellular viability, Journal of Biological Chemistry 287, 43223–43233.

40. Avila, J. (2006) Tau phosphorylation and aggregation in Alzheimer’s disease pathology, FEBS letters 580, 2922–2927.

41. Sun, L.-H., Ban, T., Liu, C.-D., Chen, Q.-X., Wang, X., Yan, M.-L., Hu, X.-L., Su, X.-L., Bao, Y.-N., Sun, L.-L., Zhao, L.-J., Pei, S.-C., Jiang, X.-M., Zong, D.-K., and Ai, J. (2015) Activation of Cdk5/p25 and tau phosphorylation following chronic brain hypoperfusion in rats involves microRNA-195 down-regulation, Journal of neurochemistry 134, 1139–1151.

42. Pan, H.-J., Wang, R.-L., Xiao, J.-L., Chang, Y.-J., Cheng, J.-Y., and Lee, C.-H. (2014) Using optical profilometry to characterize cell membrane roughness influenced by amyloid-beta 42 aggregates and electric fields, Journal of biomedical optics 19, 011009.

43. Lee, C.-W., Jang, L.-L., Pan, H.-J., Chen, Y.-R., Chen, C.-C., and Lee, C.-H. (2016) Membrane roughness as a sensitive parameter reflecting the status of neuronal cells in response to chemical and nanoparticle treatments, Journal of nanobiotechnology 14, 9.

44. Asai, H., Ikezu, S., Tsunoda, S., Medalla, M., Luebke, J., Haydar, T., Wolozin, B., Butovsky, O., Kügler, S., and Ikezu, T. (2015) Depletion of microglia and inhibition of exosome synthesis halt tau propagation, Nature neuroscience 18, 1584.

45. Lian, H., Litvinchuk, A., Chiang, A. C. A., Aithmitti, N., Jankowsky, J. L., and Zheng, H. (2016) Astrocyte-microglia cross talk through complement activation modulates amyloid pathology in mouse models of Alzheimer’s disease, Journal of Neuroscience 36, 577–589.

